# Neuropilin-1 is a host factor for SARS-CoV-2 infection

**DOI:** 10.1101/2020.06.05.134114

**Authors:** James L. Daly, Boris Simonetti, Carlos Antón-Plágaro, Maia Kavanagh Williamson, Deborah K. Shoemark, Lorena Simón-Gracia, Katja Klein, Michael Bauer, Reka Hollandi, Urs F. Greber, Peter Horvath, Richard B. Sessions, Ari Helenius, Julian A. Hiscox, Tambet Teesalu, David A. Matthews, Andrew D. Davidson, Peter J. Cullen, Yohei Yamauchi

**Author notes:** These authors contributed equally (James L. Daly, Boris Simonetti). These authors contributed equally (Carlos Antón-Plágaro, Maia Kavanagh Williamson). These authors jointly supervised this work and are joint corresponding authors.

## Abstract

SARS-CoV-2 is the causative agent of COVID-19, a coronavirus disease that has infected more than 6.6 million people and caused over 390,000 deaths worldwide^1,2^. The Spike (S) protein of the virus forms projections on the virion surface responsible for host cell attachment and penetration. This viral glycoprotein is synthesized as a precursor in infected cells and, to be active, must be cleaved to two associated polypeptides: S1 and S2^(3,4)^. For SARS-CoV-2 the cleavage is catalysed by furin, a host cell protease, which cleaves the S protein precursor at a specific sequence motif that generates a polybasic Arg-Arg-Ala-Arg (RRAR) C-terminal sequence on S1. This sequence motif conforms to the C-end rule (CendR), which means that the C-terminal sequence may allow the protein to associate with cell surface neuropilin-1 (NRP1) and neuropilin-2 (NRP2) receptors^5^. Here we demonstrate using immunoprecipitation, site-specific mutagenesis, structural modelling, and antibody blockade that, in addition to engaging the known receptor ACE2, S1 can bind to NRP1 through the canonical CendR mechanism. This interaction enhances infection by SARS-CoV-2 in cell culture. NRP1 thus serves as a host factor for SARS-CoV-2 infection, and provides a therapeutic target for COVID-19.

---

A striking difference in the S protein of SARS-CoV-2 and SARS-CoV is the presence, in the former, of a polybasic sequence motif, RRAR, at the S1/S2 boundary. It provides a cleavage site for a proprotein convertase, furin, a membrane-bound host cell protease^3–5^ (**Figure 1A**). The resulting two proteins, S1 and S2, remain non-covalently associated, with the serine protease TMPRSS2 further priming the S2 protein by proteolytic cleavage^6^. Several observations indicate that the furin-mediated processing of the S protein increases the infection and affects the tropism of SARS-CoV-2^(3–5)^. Proprotein convertase inhibitors that target furin robustly diminish SARS-CoV-2 entry into Calu-3 and HeLa cells exogenously expressing ACE2^(7)^. Moreover, furin knockdown impairs S processing, and abrogation of the polybasic site in the S reduces syncytia formation in infected cells^4,7^.

**Figure 1.**
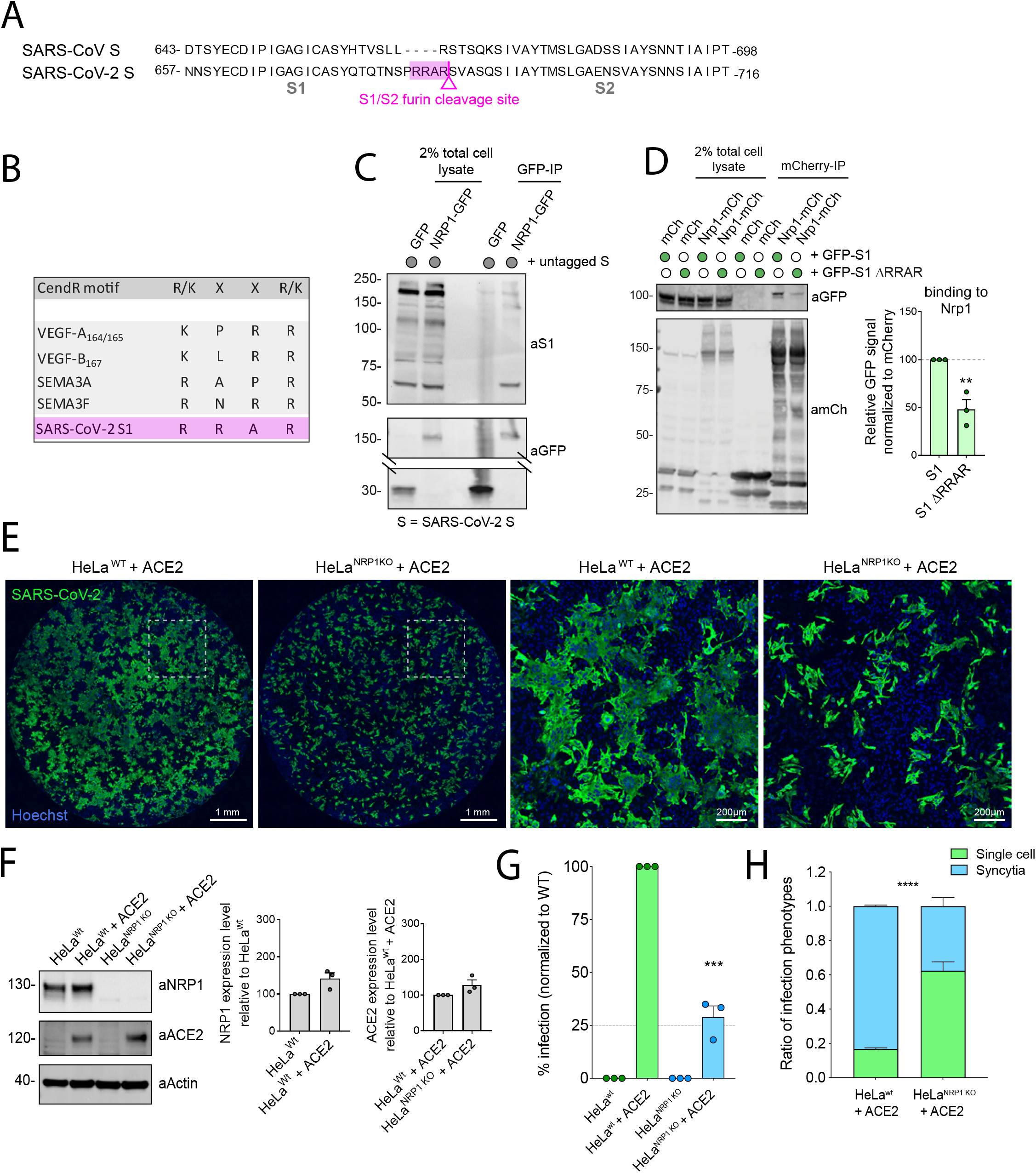
NRP1 interacts with S1 and enhances SARS-CoV-2 infection. (**A**). Alignment of the S protein sequence of SARS-CoV and SARS-CoV-2; SARS-CoV-2 S possesses a furin cleavage site at the S1/S2 boundary. (**B**). Illustration depicting the CendR motif binding to NRPs, the box highlights the similarity between well-established NRP1 ligands and the C-terminal-RRAR motif of SARS-CoV-2 S1. (**C**). Co-immunoprecipitation of the SARS-CoV-2 S protein by GFP-tagged NRP1 in HEK293T cells. HEK293T cells lentivirally transduced to express untagged SARS-CoV-2 S protein were transiently transfected with GFP or NRP1-GFP and subjected to a GFP-trap based immunoprecipitation. (**D**). CendR motif dependent co-immunoprecipitation of the SARS-CoV-2 S1 protein by Nrp1 in HEK293T cells co-transfected to express GFP-tagged S1 or GFP-S1 ΔRRAR and mCherry-tagged Nrp1. Band intensities following mCherry-nanotrap based immunoprecipitation were quantified from n=3 independent experiments using Odyssey software. The band intensities, normalized to mCherry expression, are presented as the average fraction relative to the amount of S1 protein immunoprecipitated by Nrp1-mCherry. Two-tailed unpaired t-test; P= 0.007. **(E**). Spinning-disk confocal images (20x objective, 7 z-stacks, 80 fields per well, maximum intensity projected) of HeLa^WT^ + ACE2 and HeLa^NRP1KO^ + ACE2 cells infected with SARS-CoV-2 for 16 hours. Cells were fixed and stained with anti-SARS nucleocapsid (N) and Hoechst to stain nuclei. A blow-up of the dotted square region is shown in the corresponding right-hand panel for each cell line. Images were stitched using CIDRE. (**F**). Expression of NRP1 and ACE2 in HeLa^WT^, HeLa^WT^ + ACE2, HeLa^NRP1KO^ and HeLa^NRP1KO^ + ACE2 cells. NRP1 bands were quantified from n=3 independent experiments using Odyssey software. The band intensities, normalized to Actin expression, are presented as the average fraction relative to the amount of NRP1 in HeLa^WT^. Two-tailed unpaired t-test; P= 0.0505. ACE2 bands were quantified from n=3 independent experiments using Odyssey software. The band intensities, normalized to Actin expression, are presented as the average fraction relative to the amount of ACE2 in HeLa^WT^ + ACE2. Two-tailed unpaired t-test; P= 0.1065. (**G**). SARS-CoV-2 infection in HeLa^WT^, HeLa^WT^+ ACE2, HeLa^NRP1KO^ and HeLa^NRP1KO^ + ACE2 cells. Two-tailed unpaired t-test; P= 0.0002 (n=3 independent experiments). (**H**). The ratio of single-cell and multi-cell (syncytia) infections within the infected-cell population in HeLa^WT^ + ACE2 or HeLa^NRP1KO^ + ACE2 cells were quantified. The bars, error bars and circles represent the mean, s.e.m., and individual data points. Two-tailed unpaired t-test; P<0.00001 (n=3 independent experiments). The bars, error bars and circles represent the mean,s.e.m. and individual data points, respectively. *P< 0.05, **P< 0.01, ***P< 0.001,****P< 0.0001.

We noticed that the C-terminus of the S1 protein generated by furin has a polybasic amino acid sequence (^682^RRAR^685^), that conforms to a [R/K]XX[R/K] motif, termed the ‘C-end rule’ (CendR) (**Figure 1B**)^8^. CendR motifs bind to neuropilin-1 (NRP1) and NRP2, dimeric transmembrane receptors that regulate pleiotropic biological processes, including axon guidance, angiogenesis and vascular permeability^8–10^.

To explore the possible association of the SARS-CoV-2 S1 protein with neuropilins we engineered a HEK293T cell line to stably express SARS-CoV-2 S protein. In this line we transiently expressed full length NRP1 tagged at the C-terminus with GFP (NRP1-GFP). Following a GFP-nanotrap, the isolates were probed with an antibody raised against S1. It was found that NRP1-GFP was associated with a number of proteins some of which could be S1-processed versions of the S protein **(Figure 1C)**. To test this possibility, we transiently co-expressed HEK293T cells with GFP-tagged S1 (GFP-S1) and mCherry-tagged mouse Nrp1 (Nrp1-mCherry). mCherry-nanotrap established that Nrp1 associated with the S1 protein (**Figure 1D** – comparable binding was also observed with NRP2-mCherry (**Extended Figure 1A**). That deletion of the terminal ^682^RRAR^685^ residues reduced S protein binding indicated the association largely depended on the CendR motif (**Figure 1D**). Residual binding was observed with the ΔRRAR mutant indicating an additional CendR-independent association between Nrp1 and the S1 protein.

To establish the functional relevance of this interaction, we first generated HeLa wild type and NRP1 knock out cell lines stably expressing ACE2, designated as HeLa^WT^+ACE2 and HeLa^NRP1KO^+ACE2, respectively. The two cell lines were then infected with SARS-CoV-2 (isolate SARS-CoV-2/human/Liverpool/REMRQ001/2020) and the cells fixed 16 hours post infection (h.p.i.) (**Figure 1E**). The level of ACE2 protein expression was comparable between these lines (**Figure 1F**). Infection was visualized and quantified by staining with an anti-SARS nucleocapsid (N) polyclonal antibody (**Figure 1G**). The percentage of infected cells was 73% lower in the HeLa^NRP1KO^+ACE2 cells compared to HeLa^WT^+ACE2 cells (**Figure 1G**).

We also noted that SARS-CoV-2-infected HeLa^WT^+ACE2 cells displayed a distinct multi-nucleated syncytia cell pattern, as reported by others^5^. We developed an image analysis algorithm using supervised machine learning to distinguish and quantify the multi-nucleated and single-cell infections of SARS-CoV-2 (**Extended Figure 2**)^11^. Of infected HeLa^WT^+ACE2 cells, the majority (83%) formed syncytia while in HeLa^NRP1KO^+ACE2 cells this multi-nucleated cell phenotype was less prevalent (**Figure 1H**). Together these data establish that NRP1 promotes SARS-CoV-2 infection.

**Figure 2.**
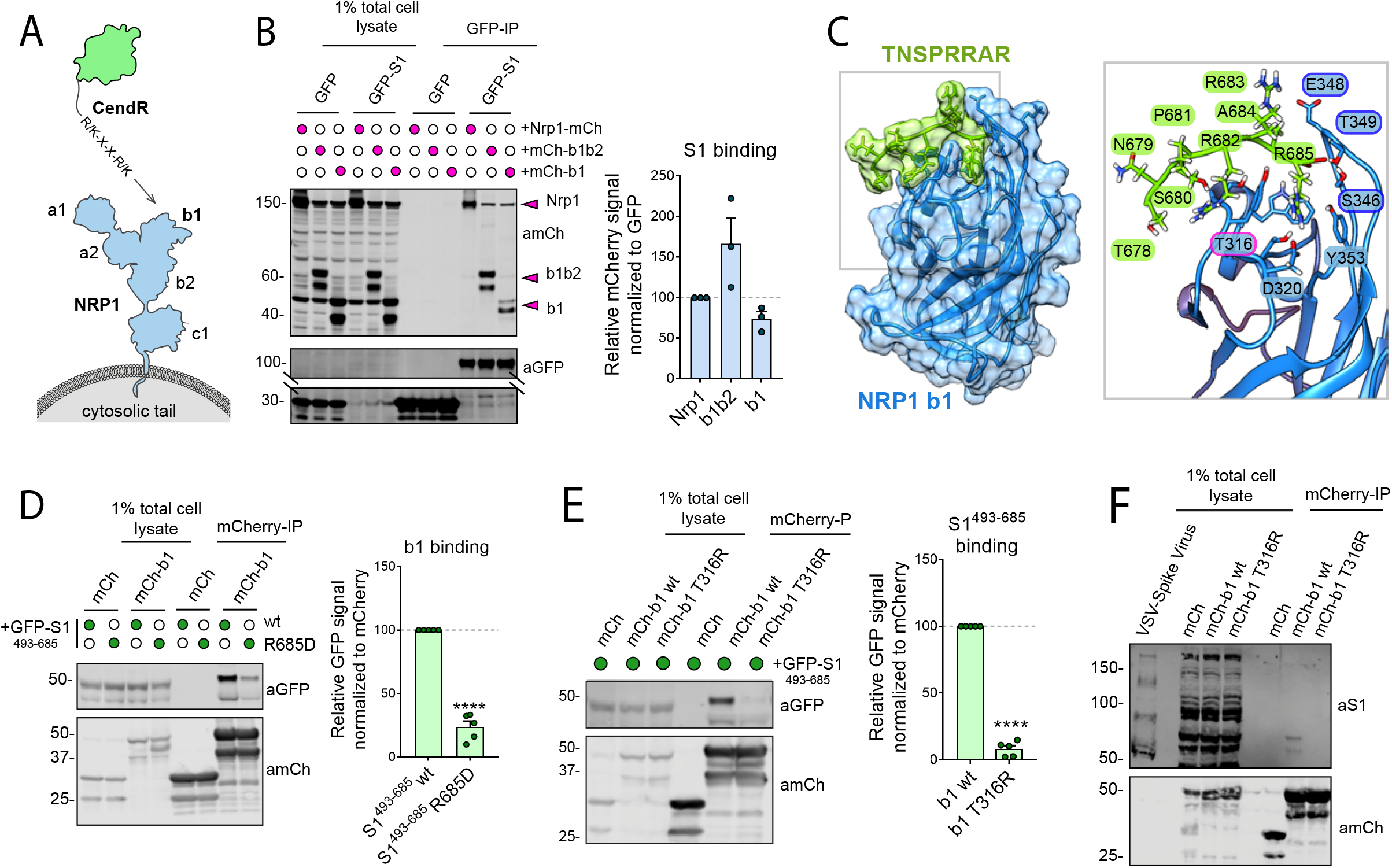
Molecular basis for CendR binding of S1 with NRP1. (**A**). Schematic of NRP1 ectodomain and CendR motif binding to b1 domain. (**B**). Co-immunoprecipitation of Nrp1 and the isolated b1b2 and b1 domains by the SARS-CoV-2 S1 protein in HEK293T cells. HEK293T cells were co-transfected with combinations of GFP or GFP-tagged S1, and mCherry-tagged Nrp1, NRP1 b1b2 or NRP1 b1 and subjected to GFP-trap based immunoprecipitation. Band intensities (highlight by arrow heads) were measured from n = 3 independent experiments using Odyssey software. Intensities were normalized to GFP expression and presented as the average fraction relative to the amount of Nrp1 immunoprecipitated by GFP-S1. One-way ANOVA and Dunnett’s test; P= 0.0845 (b1b2) and 0.5513 (b1).(**C**). Molecular models of the SARS-CoV-2 S1 C-terminus (^678^TNSPRRAR^685^), green, bound to the b1 domain of NRP1, blue, based on the crystal structure of NRP1 bound to the CendR motif of an established ligands VEGF-A164. Residue T316 is highlighted with a magenta outline as it defines the critical site for CendR binding and its corresponding T316R mutant is used later in this study. The blue outlines define the residues at the entry of the CendR binding pocket that are targeted but the mAb#3 antibody later used in this study. (**D**). Co-immunoprecipitation of SARS-CoV-2 S1^493-685^ or S1^493-685^ R685D mutant by the isolated NRP1 b1 domain in HEK293T cells. HEK293T cells were co-transfected with combinations of GFP, GFP-tagged S1^493-685^ and S1^493-685^ R685D, and mCherry or mCherry-NRP1 b1, and subjected to a mCherry-nanotrap. Band intensities were measured from n = 5 independent experiments using Odyssey software. Intensities, normalized to GFP expression, are presented as the average fraction relative to the amount of GFP-S1^493-685^ immunoprecipitated by mCherry-b1. Two-tailed unpaired t-test; <0.0001. (**E**). Co-immunoprecipitation of the SARS-CoV-2 S1^493-685^ by the isolated NRP1 b1 or b1 T316R mutant in HEK293T cells. HEK293T cells were co-transfected with combinations of GFP-tagged S1^493-685^ and mCherry, mCherry-NRP1 b1 or mCherry-NRP1 b1 T316R mutants and subjected to mCherry-nanotrap. Band intensities were measured from n = 5 independent experiments using Odyssey software. Intensities, normalized to mCherry expression, are presented as the average fraction relative to the amount of S1^493-685^ immunoprecipitated by mCherry-b1. Two-tailed unpaired t-test; P <0.0001. (**F**). Co-immunoprecipitation of the VSV pseudoparticles expressing SARS-CoV-2 S (VSV-S) by the NRP1 b1 or b1 T316R mutant isolated from HEK293T cells. HEK293T cells were co-transfected with mCherry, mCherry-NRP1 b1 or mCherry-NRP1 b1 T316R and the mCherry-tagged proteins were captured on mCherry-beads. VSV-S pseudoparticles were incubated with mCherry-beads and subjected mCherry-nanotrap (n = 2 independent experiments). The bars, error bars and circles represent the mean, s.e.m. and individual data points, respectively. *P< 0.05, **P< 0.01, ***P< 0.001, ****P< 0.0001.

The extracellular regions of NRP1 and NRP2 are composed of two complementary binding CUB domains (a1 and a2), two coagulation factor domains (b1 and b2), and a MAM domain (c1). Of these, particularly important is the b1 domain that contains the specific binding site for CendR motifs, with the b2 domain further stabilising this interaction (**Figure 2A**)^12^. When we co-expressed the mCherry-b1 domain of human NRP1 together with GFP-S1, the single domain alone was sufficient for immunoprecipitation by the viral protein (**Figure 2B**). A construct with both b1 and b2 (mCherry-b1b2) displayed elevated association compared to b1 alone and full-length Nrp1-mCherry (**Figure 2B**). The S1 C-terminal residues 493-685 immunoprecipitated mCherry-b1b2 to a greater extent than full length S1, further narrowing the binding interface between the S1 C-terminus and NRP1 b1b2 domains (**Extended Figure 1B)**. Based on the crystal structure of NRP1 bound to the CendR motif of VEGF-A164(12), we generated molecular models of the SARS-CoV-2 S1 C-terminus (^678^TNSPRRAR^685^) bound to the b1 domain of NRP1 (**Figure 2C**). The S1 C-terminus model closely resembled that of endogenous CendR motifs bound to NRP1, consistent with the guanidino group of S1 R685 establishing a salt-bridge with the NRP1 carboxylate of D320^(12)^. In the NRP1 b1 domain, T316 sits at the base of the CendR binding pocket, which in the case of VEGF-A164 controls ligand occupancy12. The S1 C-terminal carboxylate was predicted to interact mainly with the backbone amides of NRP1, namely K347 and E348 (**Figure 2C**).

Based on this model, we investigated whether two key residues (R685 of S1 and T316 of NRP1) contributed to the interaction between S1 and NRP1. Site-directed mutagenesis of R685 of S1^493-685^ to aspartic acid reduced S1 association with the NRP1 b1 domain by greater than 75 % (**Figure 2D**). Site-directed mutagenesis of T316 to arginine within mCherry-b1 reduced association with GFP-S1^493-685^, more than 80%, consistent with its inhibitory impact on VEGF-A164 binding12 (**Figure 2E)**. Finally, to evaluate the interaction of NRP1 with trimeric S on the surface of virions, we incubated vesicular stomatitis virus (VSV) pseudoviral particles harbouring SARS-CoV-2 S protein with wild type mCherry-b1 or mCherry-b1(T316R) immobilised on mCherry-nanotrap beads (**Figure 2F**). Wild type mCherry-b1 immunoprecipitated proteins that could correspond to processed forms of S1, whereas the T316R mutant did not (**Figure 2F**).

To test the functional relevance of this association, we transiently expressed GFP, NRP1-GFP or NRP1(T316R)-GFP in HeLa^NRP1KO^+ACE2 cells. Wild type and NRP1(T316R) mutant expression and ACE2 expression levels were comparable (**Figure 3A**) and like wild type NRP1, NRP1(T316R)-GFP retained a localisation to the cell surface (**Figure 3B**). When these cells were infected with SARS-CoV-2, viral infection was significantly enhanced in cells expressing NRP1-GFP compared to GFP control, whereas cells expressing the T316R mutant failed to rescue infection (**Figure 3C**). This established that the SARS-CoV-2 S1 CendR (^682^RRAR^685^) and NRP1 interaction promotes infection.

**Figure 3.**
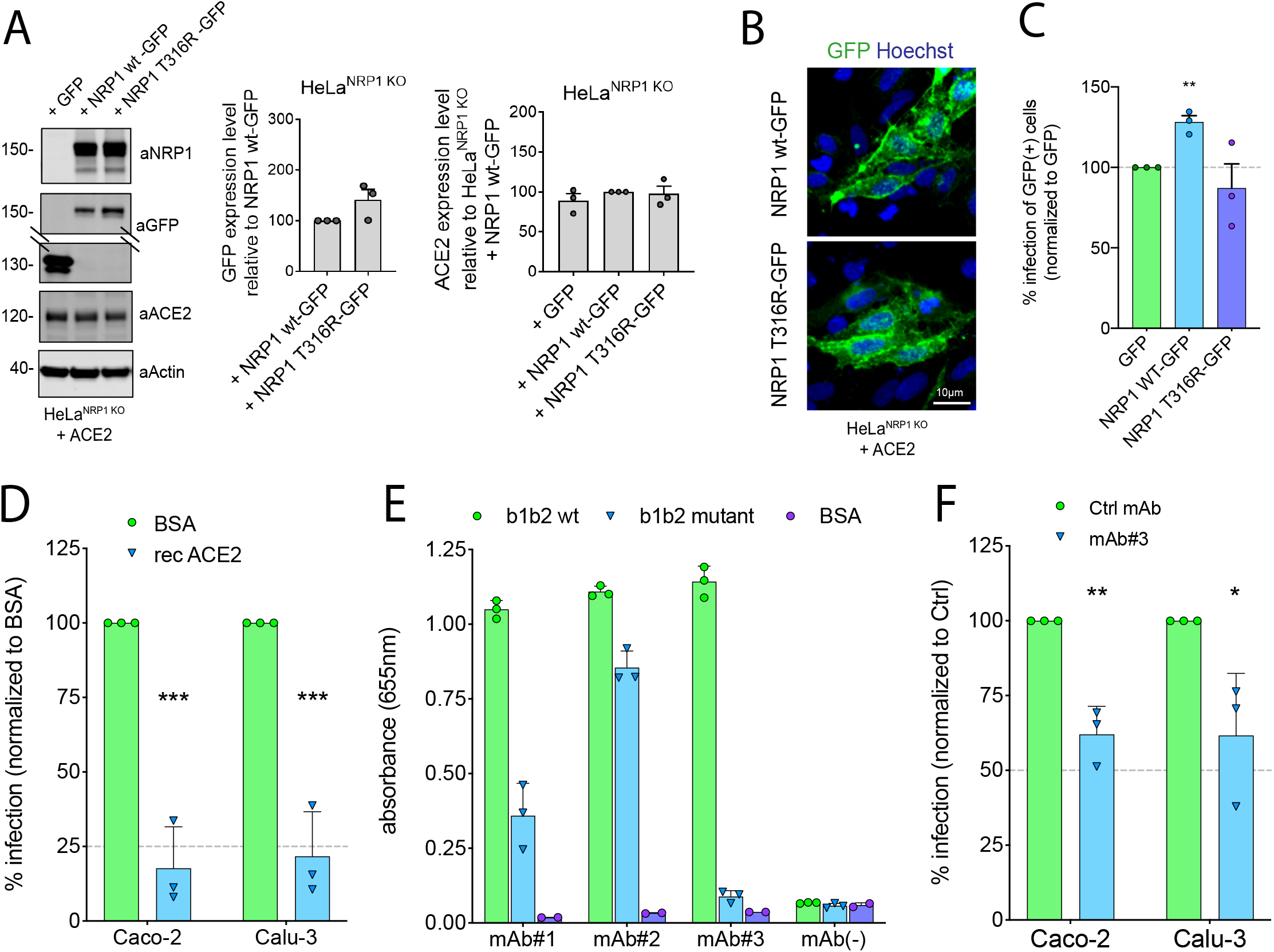
CendR is required for SARS-CoV-2 infection. (**A**). HeLa^NRP1KO^ + ACE2 cells transiently expressing NRP1-GFP or NRP1(T316R)-GFP were harvested and analysed by western blot. Band intensities were measured from n=3 independent experiments using Odyssey software. GFP bands were quantified from n=3 independent experiments using Odyssey software. The band intensities, normalized to Actin expression, are presented as the average fraction relative to the amount of NRP1 wt-GFP. Two-tailed unpaired t-test; P=0.1167. ACE2 bands were quantified from n=3 independent experiments using Odyssey software. The band intensities, normalized to Actin expression, are presented as the average fraction relative to the amount of ACE2 in HeLa^NRP1KO^ + NRP1 wt-GFP. one-way analysis of variance (ANOVA) and Dunnett’s test; P= 0.5293 (HeLa^NRP1KO^ + GFP), P= 0.9672 (HeLa^NRP1KO^ + NRP1 T316R-GFP) (**B**). Spinning-disk confocal images of HeLa^NRP1KO^+ ACE2 cells expressing NRP1-GFP or NRP1(T316R)-GFP. Hoechst was used to stain nuclei (blue). Maximum projection images of z-stacks acquired with a 20x objective using an automated spinning-disk confocal microscope. Scale bar; 10μm. **(C).** HeLa^NRP1KO^ + ACE2 cells transfected with GFP, NRP1-GFP or NRP1(T316R)-GFP constructs were infected 24 h later with SARS-CoV-2. At 16 h.p.i. the cells were fixed and stained for SARS-CoV-2-N, and viral infection quantified in the GFP-positive subpopulation of cells. The percentage of infection was normalized to that of GFP-transfected cells. Two-tailed unpaired t-test; p=0.002 (n=3 independent experiments) **(D).** Inhibition of SARS-CoV-2 infection by treatment with recombinant ACE2 in Caco-2 and Calu-3 cells. Cells were pre-treated with recombinant ACE2 (10 μg/mL) for 1 h prior to SARS-CoV-2 infection. At 16 h.p.i. the cells were fixed and stained for SARS-CoV-2-N, and infection was quantified. Two-tailed unpaired t-test; P <0.001 (n=3 independent experiments) (**E**). ELISA of anti-NRP1 monoclonal antibodies (mAb#1, mAb#2, mAb#3) diluted 1/10 using plates coated with NRP1 b1b2 wild type, b1b2 mutant (S346A, E348A, T349A) or BSA, used as control. No mAb was also used as a control. Binding is represented as arbitrary units of absorbance at 655 nm. (**F**). mAb#3 reduces SARS-CoV-2 infection in Caco-2 and Calu-3 cells. Cells were pretreated with 50 μg/mL of anti-avian influenza A virus hemagglutinin (H11N3) (Ctrl) mAb or mAb#3 for 1 h followed by infection with SARS-CoV-2. Cells were fixed at 16 h.p.i. and stained for SARS-CoV-2-N and Hoechst to stain nuclei. p=0.002 (Caco-2) and p=0.032 (Calu-3) (n=3 independent experiments). The bars, error bars and circles represent the mean, s.e.m. and individual data points, respectively. *P<0.05, **P<0.01, ***P<0.001, ****P<0.0001.

To evaluate the biological importance of the SARS-CoV-2 S1 interaction with NRP1, we turned to analysing infection of Caco-2 (human colon adenocarcinoma) and Calu-3(human lung cancer) cells. Incubation of these cells with recombinant ACE2 inhibited SARS-CoV-2 infection by greater than 75%, presumably by binding to S1 and inhibiting interaction with ACE2 on the cell surface (**Figure 3D**). To establish the importance of the interaction with NRP1, we screened a series of monoclonal antibodies (mAb#1, mAb#2, mAb#3) raised against the NRP1 b1b2 ectodomain. All three monoclonal antibodies bound to the NRP1 b1b2 domain but only mAb#3 bound to the CendR-binding pocket (**Figure 3E, Extended Figure 3A**), as defined by a reduced ability to bind to a b1b2 mutant that targets residues (S346, E348, T349) at the opening of the binding pocket^12^ (**Figure 2C**). All three monoclonal antibodies displayed staining by immunofluorescence in NRP1-expressing PPC-1 human primary prostate cancer cells but not in M21 human melanoma cells that do not express NRP1^(8)^ (**Extended Figure 3B)**. Using mAb#3 and a control monoclonal antibody targeting avian influenza A virus (H11N3) hemagglutinin we found that mAb#3 inhibited SARS-CoV-2 infection of Caco-2 and Calu-3 cells by 38% (**Figure 3F**).

The molecular and cell biological mechanisms of SARS-CoV-2 infection remain largely unexplored. Cell entry of SARS-CoV-2 depends on priming by host cell proteases, including at the S1/S2 site, and the S2’ site which drives fusion with cellular membranes^5,6,14^. Our data indicate that the SARS-CoV-2 S-protein binds to cell surface receptor NRP1 via the S1 CendR motif generated by the furin cleavage of S1/S2. This interaction promotes infection by SARS-CoV-2 in physiologically relevant cell lines widely used in the study of COVID-19. The molecular basis for the effect is unclear, but neuropilins are known to mediate the internalisation of CendR ligands through an endocytic process resembling macropinocytosis, and additionally serve as activators of receptor tyrosine kinases to mediate cell signalling^8,15^. S1 binding to NRP1 may therefore contribute to viral entry and survival within the host cell. Interstingly, gene expression analysis of lung tissue from COVID-19 patients recently revealed an up-regulation of NRP1 and NRP2^(16)^.

A SARS-CoV-2 virus with a natural deletion of the S1/S2 furin cleavage site demonstrated attenuated pathogenicity in hamster models, causing less alveolar damage^17^. NRP1 binding to the CendR-motif in S1 is thus likely to play a role in the increased pathogenicity of SARS-CoV-2 compared with SARS-CoV. The interaction between S1 and NRP1 can be reduced by monoclonal antibodies that bind to the CendR-binding pocket of NRP1 b1. Disrupting CendR-peptide binding to neuropilins by antibodies and oligopeptide/peptidomimetics has been recognised as a potential anti-cancer strategy^18–25^, and the same approach may provide a route to COVID-19 therapies.

It is noteworthy that many other viruses including highly pathogenic influenza, alpha-, flavi- and retroviruses undergo furin cleavages and acquire a C-terminus that conforms to the CendR, which make them potential ligands for neuropilin binding^26–28^. They also include certain human β-coronaviruses of separate lineages such as Middle Eastern respiratory syndrome coronavirus (MERS)-CoV, and human coronavirus HKU1 (HCoV-HKU1)^(28)^. Taken together, we conclude that neuropilins are emerging targets in the urgent research and treatment of COVID-19 and, more broadly, important host factors to be considered in the wider context of viral entry mechanisms.

## MATERIALS AND METHODS

### Antibodies and reagents

The following antibodies were used in this study: mouse anti-β actin (Sigma-Aldrich, A1978, WB 1:2000), mouse anti-ACE2 (Proteintech, 66699-1-Ig, WB 1:1000), mouse anti-GFP (Roche, 11814460001, WB 1:2000), rabbit anti-mCherry (Abcam, ab167453, WB 1:2000), rabbit anti-NRP1 (Abcam, ab81321, WB 1:1000), rabbit anti-SARS-CoV-2 Spike RBD (S1 epitope) (Sino Biologicals, 40592-T62, WB 1:1000), rabbit anti-SARS nucleocapsid (N) polyclonal antibody (ROCKLAND, 200-401-A50, IFA 1:2000), mouse anti-SARS-CoV-2 Spike antibody [1A9] (S2 epitope) (GeneTex, GTX632604, WB 1:1000). Recombinant human ACE2 was purchased from Sino Biological (10108-H08H). The monoclonal antibody against the hemagglutinin of influenza A/duck/New Zealand/164/76(H11N3) was a kind gift of Robert Webster.

### Cell culture and transfection

Calu-3, Caco-2 (a kind gift from Dr Darryl Hill), HeLa, HEK293T and Vero E6 cell lines were originally sourced from the American Type Culture Collection. Authentication was from the American Type Culture Collection. We did not independently authenticate the cell lines. Cells were grown in DMEM medium (Sigma-Aldrich) supplemented with 10% (vol/vol) FCS (Sigma-Aldrich) and penicillin/streptomycin (Gibco) with the exception of Calu-3 cells that were grown in Eagle’s minimal essential medium (MEM+GlutaMAX; Gibco™, ThermoFischer) supplemented with 10% FCS 0.1mM non-essential amino acids (NEAA), 1mM sodium pyruvate, 100 IU/ml streptomycin and 100 μg/ml penicillin. Caco-2 cells were maintained in DMEM+GlutaMAX, 10% FCS and 0.1mM NEAA. FuGENE HD (Promega) was used for transient transfection of DNA constructs for infection assays according to the manufacturer’s instructions.

PPC-1 human primary prostate cancer cells were obtained from Erkki Ruoslahti laboratory at Cancer Research Center, Sanford-Burnham-Prebys Medical Discovery Institute. M21 human melanoma cells were obtained from David Cheresh at University of California San Diego. Cells were cultured in DMEM medium containing 100 IU/mL of streptomycin, penicillin, and 10% FBS in 37°C incubator with 5% CO2.

To generate a NRP1-null HeLa cell line, the following guide RNA was cloned into pSpCas9(BB)-2A-Puro (PX459): 5’-GATCGACGTTAGCTCCAACG-3’. gRNA was transfected into HeLa cells using FuGENE HD. 24 hours later, transfected cells were selected with puromycin. Selected cells were trypsinised and diluted to a concentration of 2.5 cells mL^−1^ in Iscove’s modified Dulbecco’s medium (Gibco) supplemented with 10% (vol/vol) FBS (Sigma-Aldrich). 200 μL of this suspension was plated into 96-well plates to seed single cell colonies. After three weeks, colonies were expanded and lysed, and knockout was validated by immunoblotting for NRP1.

### SARS-CoV-2 isolation and infection

A clinical specimen in viral transport medium, confirmed SARS-CoV-2 positive by qRT-PCR (kindly proved by Dr Lance Turtle, University of Liverpool), was adjusted to 2 ml with OptiMEM (Gibco™, ThermoFisher), filtered through a 0.2 μm filter and used to infect Vero E6 cells. After 1 h the inoculum was diluted 1:3 (vol/vol) with MEM supplemented with 2% FCS and incubated at 37 °C in a 5% CO2 incubator for 5 days. The culture supernatant was passaged twice more on Vero E6 cells until cytopathic effect was observed and then once on Caco-2 cells to produce the stock used in the experiments. The intracellular viral genome sequence and the titre of virus in the supernatant were determined as previously described^29^ and the virus termed SARS-CoV-2/human/Liverpool/REMRQ0001/2020. For virus infections, virus was added directly to culture medium of the target cells in a 96-well plate at the required infectious dose and the plates incubated at 37 °C for 16 h. The culture supernatant was removed, and the cells fixed with 4% (vol/vol) paraformaldehyde (PFA) for 1 h at room temperature. All work with infectious SARS-CoV-2 was done inside a class III microbiological safety cabinet in a containment level 3 facility at the University of Bristol.

### Pseudotyped virus generation

The VSVΔG system was used to generate pseudovirus particles decorated with SARS-2-S as described previously^5,30^ with some modifications. Briefly, HEK293T cells were grown in 100 mm diameter dishes to 90% confluency and were subsequently transfected with 6 μg of pCG1_SARS-2-S plasmid using polyethylenimine (PEI) Max (MW = 40,000KDa, Polysciences, Germany) as transfection reagent. Transfection was performed using a PEI:DNA ratio of 4:1 in serum free DMEM for 4 hours at 37°C. The cells were then washed with PBS and cultured in fresh complete DMEM supplemented with 5% FBS at 37°C overnight. The next day cells were exposed to the replication deficient VSV*ΔG-fLuc vector (kindly provided by Markus Hoffmann, German Primate Center, Leibnitz) for 2 hours at 37°C. The cells were then washed with PBS before the addition of medium supplemented with anti-VSV-G I1 antibody (kindly provided by Markus Hoffmann, German Primate Center, Leibnitz). The cells were further incubated at 37°C for 24 hours before the supernatant was harvested and clarified by centrifugation at 2,000×g for 10 minutes. For immunoprecipitation experiments, VSV pseudoviral particles were concentrated 10-fold using 100KDa Amicon^®^ Ultra centrifugal filter units.

### Infection assays, indirect immunofluorescence, automated confocal imaging

For infection assays, cells seeded in Clear 96-well Microplates (Greiner Bio-one) were infected with SARS-CoV-2/human/Liverpool/REMRQ001/2020 isolate in MEM, 2% FCS, supplemented with 0.1mM NEAA and fixed in 4% PFA in PBS at 16 h.p.i. After permeabilisation with 0.1% TritonX-100 in PBS, 1% BSA, the cells were blocked and stained in 1% BSA in PBS containing anti-SARS Nucleocapsid (N) rabbit polyclonal antibody or anti-SARS Spike (S) monoclonal antibody and further stained with Hoechst (1:10000) and appropriate Alexa Fluor (488/594/647)-conjugated secondary antibodies (Thermo Fisher Scientific). The stained plates were imaged using an automated high-content spinning-disk microscope CQ1 (Confocal Quantitative Image Cytometer, Yokogawa, Japan) using UPlanSApo 10x/0.4na, UPlanSApo 20x/0.75na or UPlanSApo 40x/0.95na objectives (Olympus, Japan). To capture a single 96-well, 20 fields (with 10x objective) or 80 fields (20x) were imaged by z-stacks at 5 μm intervals and maximum intensity projected for analysis. Yokogawa CQ1 imaging was performed with four excitation laser lines (405/488/561/640nms) with spinning disc confocal.

### Cell immunostaining and confocal microscopy

PPC-1 and M21 cells were seeded in a 24-well plate (50,000 cells/well) with coverslips and allowed to grow until the next day. The medium was removed, the cells were washed twice with PBS pH 7.4, and fixed with 4% PFA in PBS for 10 min at room temperature. The cells were washed twice with PBS and once with PBST (PBS with 0.05 % Tween-20). Blocking buffer (5% BSA, 5% FBS, 5% goat serum in PBST) was added to the cells and incubated for 1 h at room temperature. The blocking buffer was removed and 0.3 mL of monoclonal antibodies were added to the cells (mAB diluted 1 in 5 in diluted blocking buffer containing 1% BSA, 1% FBS, 1% goat serum in PBST). Cells were incubated for 1 h at room temperature and washed 3 times with PBST. Secondary antibody AlexaFluor 546 goat anti-mouse IgG (Invitrogen Molecular Probes, Cat. No. A11003) was added to the cells (4μg/mL in diluted blocking buffer). Cells were incubated for 30 min at room temperature, washed 3 times with PBS and stained with 1μg/mL of DAPI for 10 min at room temperature. After three washes with PBS, the cells were mounted on glass slides with mounting media (Fluoromount-G; Electron Microscopy Sciences) and sealed with nail polish. A confocal microscope FV1200MPE (Olympus, Japan) was used for cell imaging with an UPlanSApo 60x/1.35na objective (Olympus, Japan). The images were analyzed using Olympus FluoView Ver.4.2a Viewer software.

### Image analysis

Projected images taken with a 20x objective were used for image analysis for single-cell and multinucleated cell infection image analysis with supervised machine learning. Image processing was performed using the BIAS software (Single-Cell Technologies Inc., Hungary). Firstly, images of each fluorescence channel were corrected using the CIDRE illumination correction method^31^. Individual cell nuclei were detected by a deep machine learning-based segmentation algorithm NucleAIzer^11^. Cellular cytoplasm were detected both on the green and red channels using UNET to enhance fluorescence images^32^. The method was trained to precisely delineate often faint signals in the cytoplasm. Cellular phenotypes were assigned to each individual nucleus. These are infected cells which contain a single nucleus (Single Cell Infection), those that contain more multiple nuclei (Multi Nuclei Infection) as observed in the distinct cell-cell fusion syncytia phenotype. Supervised machine learning was used for phenotypic assignment. The decisions were based on single-cell and its microenvironment’s morphology and intensity features^33^. Final statistics include the number of multi-nucleated cells, the average number of nuclei in these cells and the count of other phenotypic classes. Yokogawa CQ1 was also used for image quantification.

### Generation of Stable Lentiviral Cell Lines

The genes of interest were subcloned into the lentiviral vector pLVX for the generation of lentiviral particles. Lentiviral particles were produced and harvested in HEK293T cells. HeLa cells were transduced with lentiviral particles to produce stably expressing cell lines. Transduced HeLa and HEK293T cells were grown in DMEM supplemented with 10% (vol/vol) FCS and penicillin/streptomycin and grown at 37 °C in a 5% CO2 incubator. Following transduction, pLVX-expressing cells were selected with puromycin or blasticidin accordingly.

### Immunoprecipitation and Quantitative Western Blot Analysis

Cells were lysed in PBS with 1% Triton X-100 and protease inhibitor cocktail for western blotting. The protein concentration was determined using a BCA assay kit (Thermo Fisher Scientific) and equal amounts were resolved on NuPAGE 4–12% precast gels (Invitrogen). Blotting was performed onto polyvinylidene fluoride membranes (Immobilon-FL, EMD Millipore), followed by detection using the Odyssey infrared scanning system (LI-COR Biosciences). When using the Odyssey, we routinely performed western blot analysis where a single blot was simultaneously probed with antibodies against two proteins of interest (distinct antibody species), followed by visualization with the corresponding secondary antibodies conjugated to distinct spectral dyes.

For the GFP- and mCherry-based immunoprecipitations, HEK293T cells were transfected with GFP or mCherry constructs using polyethylenimine (Sigma-Aldrich). The cells were lysed in immunoprecipitation buffer (50mM Tris–HCl, 0.5% NP-40 PBS with protease inhibitor cocktail (Roche)) 24hr after transfection and subjected to GFP-trap (ChromoTek) or RFP-tap (ChromoTek) beads. Following immunoprecipitation, the beads were washed twice in 50mM Tris-HCl, 0.25% NP40 PBS with protease inhibitor cocktail, pH 7.5, and once in 50mM Tris-HCl PBS with protease inhibitor cocktail, pH 7.5, before boiling in 2X LDS sample loading buffer for elution. Immunoblotting was performed using standard procedures. Detection was performed on an Odyssey infrared scanning system (LI-COR Biosciences) using fluorescently labelled secondary antibodies.

For immunoprecipitation of the VSV-Spike pseudotyped virus, mCherry, mCherry-b1 and mCherry-b1 T316R were transfected into HEK293T cells the day before immunoprecipitation. Cells were lysed in 50mM Tris-HCl 0.5% NP40 PBS with protease inhibitor cocktail, pH 7.5. Lysates were cleared by centrifugation at 20,000 g for 10 minutes at 4°C. From the resulting supernatant, an input fraction was reserved, and the rest incubated with mCherry-nanotrap beads to rotate for 1 hour at 4°C. Following enrichment of mCherry constructs, the beads were washed twice in 50mM Tris-HCl 0.25% NP40 PBS with protease inhibitor cocktail, pH 7.5, and twice in 50mM Tris-HCl PBS with protease inhibitor cocktail, pH 7.5, to removed residual cell lysate and detergent. VSV-Spike pseudotyped virus was added to the isolated mCherry beads and incubated rotating for a further 1 hour at 4°C. Following virus immunoprecipitation, the beads were again washed twice in 50mM Tris-HCl 0.25% NP40 PBS with protease inhibitor cocktail, pH 7.5, and twice in 50mM Tris-HCl PBS with protease inhibitor cocktail, pH 7.5, before boiling in 2X LDS sample loading buffer for elution.

### ELISA assay with monoclonal antibodies

High affinity protein-binding 96-well plates (Nunc™ Maxisorp™ Cat No. 442404) were coated with 1 μg of protein (100 μL of 10 μg/mL of protein solution in PBS) overnight at 37°C. The wells were washed 5 times with PBS and blocked for 1 h at 37°C with blocking buffer (1% BSA, 0.1%Tween-20 in PBS). The mAb dilutions in blocking buffer were added to the wells and incubated for 1 h at 37°C. The wells were washed 5 times with blocking buffer and the peroxidase-conjugated affinity pure donkey anti-mouse IgG (Immuno Research Laboratories) was added (diluted 1 in 20.000 in blocking buffer). The plate was incubated for 1 h at 37°C and washed 5 times with blocking buffer. The peroxidase substrate (TMB Peroxidase EIA Substrate Kit #1721067, Bio-Rad) was added as described in the manufacturer instructions. The absorbance of the samples was read at 655 nm using Tecan Sunrise microplate reader (Tecan, Switzerland).

### Plasmids

The SARS-CoV-2 S gene was cloned into pLVX vectors using a commercially synthesized EGFP-S gene fusion plasmid (the S gene sequence was that of SARS-CoV-2 isolate Wuhan-Hu-1; GenBank: MN908947.3; GeneArt, ThermoFischer) as a template by Gibson assembly (NEB). For the untagged version, in brief, the S gene was amplified using overlapping primers and cloned into a ‘pLVX-MCS-T2A-Puro’ vector previously digested with EcoRI/BamHI. The isolated S1 constructs and S1 truncations were amplified from commercially synthesized plasmids (GeneArt, ThermoFischer) encompassing nucleotides 20021 – 22960 and 22891 – 28830 of the SARS-CoV-2 isolate Wuhan-Hu-1 genome (GenBank: MN908947.3) and cloned in pEGFP.C1 using KpnI/BamHI. Mouse Nrp1-mCherry was a kind gift from Donatella Valdembri. pEGFP-NRP1 and pEGFP-NRP2 were kind gifts from Mu-Sheng Zeng.

### Molecular Modelling of NRP1 b1 Domain Binding to S1 TNSPRRAR Peptide

The 4DEQ.pdb crystal structure of the VEGF-A peptide attached to the type C1 domain of NRP1 via a linker was used to model the potential for the furin-cleaved S protein from SARS-CoV-2 to bind. A model of the S protein with loops intact had been previously assembled from the cryo-EM structures pdbs 6VXX and 6VSB. From this model, the TNSPRRAR residues pertaining to the post-furin cleaved, C-terminus of the S1 domain of the S protein from SARS-CoV2 were placed into the NRP1 binding site, guided by the position of the VEGF-A PRR residues in 4DEQ.pdb.

#### System setup

Gromacs was used to prepare and run the simulation. Hydrogen atoms were added to the NRP1/S peptide complex consistent with pH 7 and surrounded by a box 2 nm larger than the polypeptide in each dimension and filled with TIP3P water. Random water molecules were replaced by sodium and chloride ions to give a neutral (uncharged overall) box and an ionic strength of 0.15 M. The system was subjected to 1000 steps of energy minimisation prior to a position-restrained trajectory to produce velocities in which the protein atoms were restrained, but the water molecules allowed to move.

#### Simulation details

The resulting system was subjected to a short molecular dynamics simulation to see if the peptide remained bound and to allow the peptide to adopt a comfortable binding position. This simulation was performed as an NPT ensemble at 310 K using periodic boundary conditions. Short range electrostatic and van der Waals’ interactions were truncated at 1.4 nm while long range electrostatics were treated with the particle-mesh Ewald’s method and a long-range dispersion correction applied. Pressure was controlled by the Berendsen barostat and temperature by the V-rescale thermostat. The leap-frog algorithm was used to integrate over a 2 fs time step, constraining bond vibrations with the P-LINCS method. Structures were saved every 0.1 ns for analysis and run over 20 ns. Simulation data were accumulated on the Bristol BrisSynBio supercomputer Bluegem.

#### Software

The GROMACS-2019.1 suite of software was used to set up and perform the molecular dynamics simulations. Molecular graphics manipulations and visualisations were performed using VMD-1.9.3 and Chimera64-1.14.

The peptide remained bound throughout and appeared to settle into an eminently plausible binding pose. There was room to easily accommodate the rest of the S1 spike protein domain when aligned with the TNSPRRAR peptide in complex with NRP1.

## ACKNOWLEDGEMENTS

JLD was supported by a Wellcome Trust studentship from the Dynamic Molecular Cell Biology Ph.D. programme (203959/Z/16/Z), CAP was supported by Beca Fundación Ramón Areces Estudios Postdoctorales en el Extranjero and MKW was supported by a MRC grant (MR/R020566/1) awarded to ADD. This project has received funding from the MRC (MR/P018807/1), Wellcome Trust (104568/Z/14/2), Lister Institute of Preventive Medicine, and Elizabeth Blackwell Institute for Health Research Rapid Response Call (COVID-19) awarded to PJC, the European Research Council under the European Union’s Horizon 2020 research and innovation programme (No 856581 - CHUbVi), and from MRC-AMED (MR/T028769/1) awarded to YY, and the United States Food and Drug Administration grant number HHSF223201510104C ‘Ebola Virus Disease: correlates of protection, determinants of outcome and clinical management’ amended to incorporate urgent COVID-19 studies awarded to JAH, ADD and DAM. We thank the Bristol Synthetic Biology Centre and the Advanced Computing Research Centre for provision of HPC (Bluegem). RH and PH acknowledge support from the LENDULET-BIOMAG Grant (2018-342), from H2020-discovAIR (874656), and from Chan Zuckerberg Initiative, Seed Networks for the HCA-DVP. TT was supported by the European Regional Development Fund (Project No. 2014-2020.4.01.15-0012), by European Research Council grant GLIOGUIDE and Estonian Research Council (grants PRG230 and EAG79, to T.T.).

## AUTHOR CONTRIBUTIONS

JLD, BS, AH, PJC and YY conceived the study. JLD, BS and YY performed most of the experiments. MKW, DAM and ADD performed all work with infectious SARS-CoV-2 supervised by ADD. CAP, MB, LSG, UFG, KK, RBS, DKS, JAH and TT did experimental work and/or provided essential reagents. RH and PH performed image analysis. BS, ADD, PJC and YY supervised the research. JLD, BS, ADD, PJC and YY wrote the manuscript and made the figures. All authors read and approved the final manuscript.

## ADDITIONAL INFORMATION

**Extended Figure 1.**
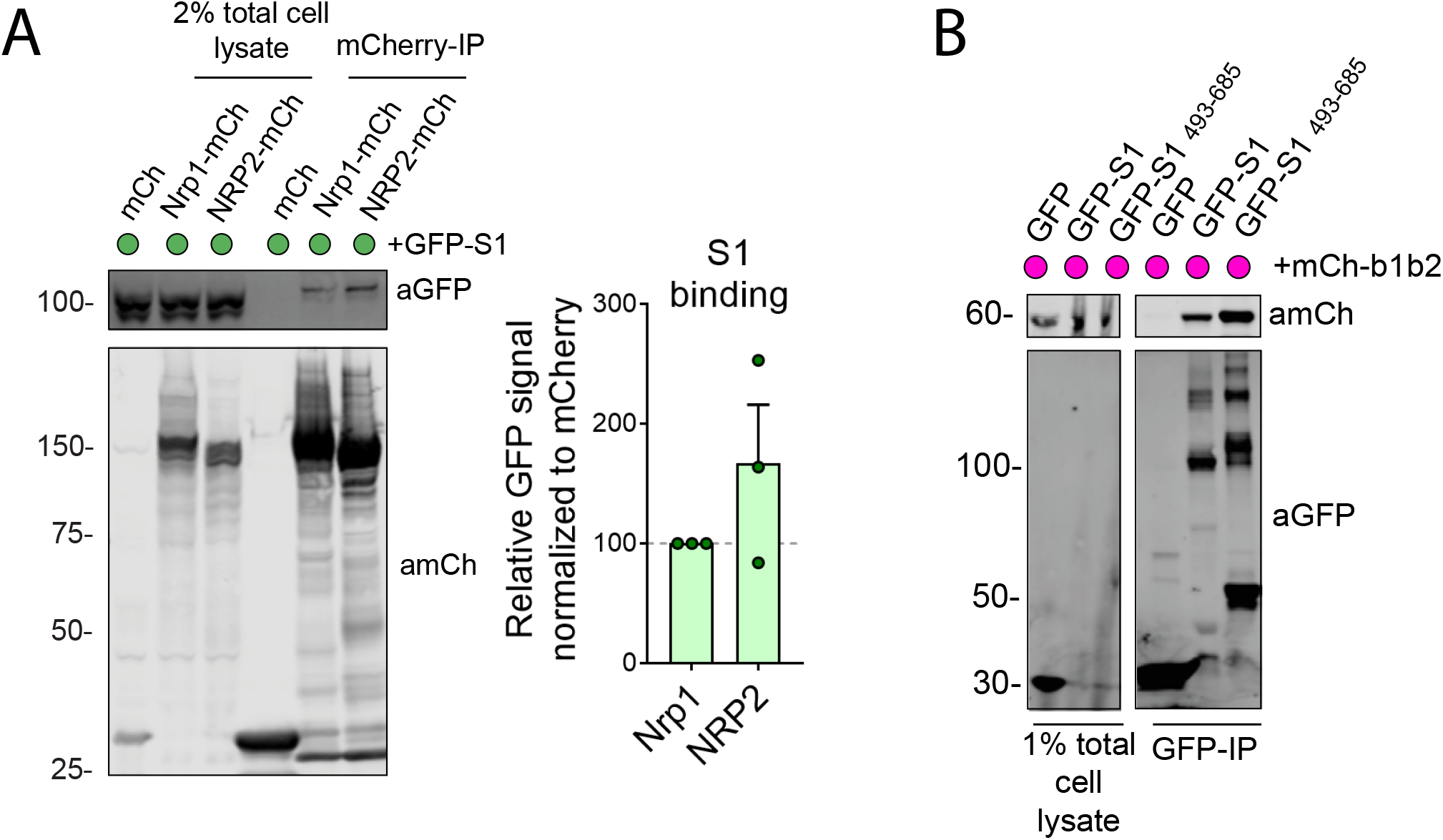
a. SARS-CoV-2 S1 protein also interacts with NRP2. Co-immunoprecipitation of the SARS-CoV-2 S1 protein by Nrp1 and NRP2 in HEK293T cells. HEK293T cells were co-transfected with combinations of GFP-tagged S1 and mCherry-tagged Nrp1 or NRP2 and subjected to a mCherry-trap based immunoprecipitation. Summary of the relative levels of binding. The band intensities were measured from n= 3 independent experiments using Odyssey software. The band intensities, normalized to mCherry expression, are presented as the average fraction relative to the amount of S1 immunoprecipitated by Nrp1-mCherry. two-tailed unpaired t-test; P= 0.2421. b. The 493-685 portion of the SARS-CoV-2 S1 protein has enhanced interaction with NRP1 b1b2. HEK293T cells were co-transfected to express GFP-tagged S1 or GFP-tagged S1^493-685^ and mCherry-tagged NRP1 b1b2.

**Extended Figure 2.**
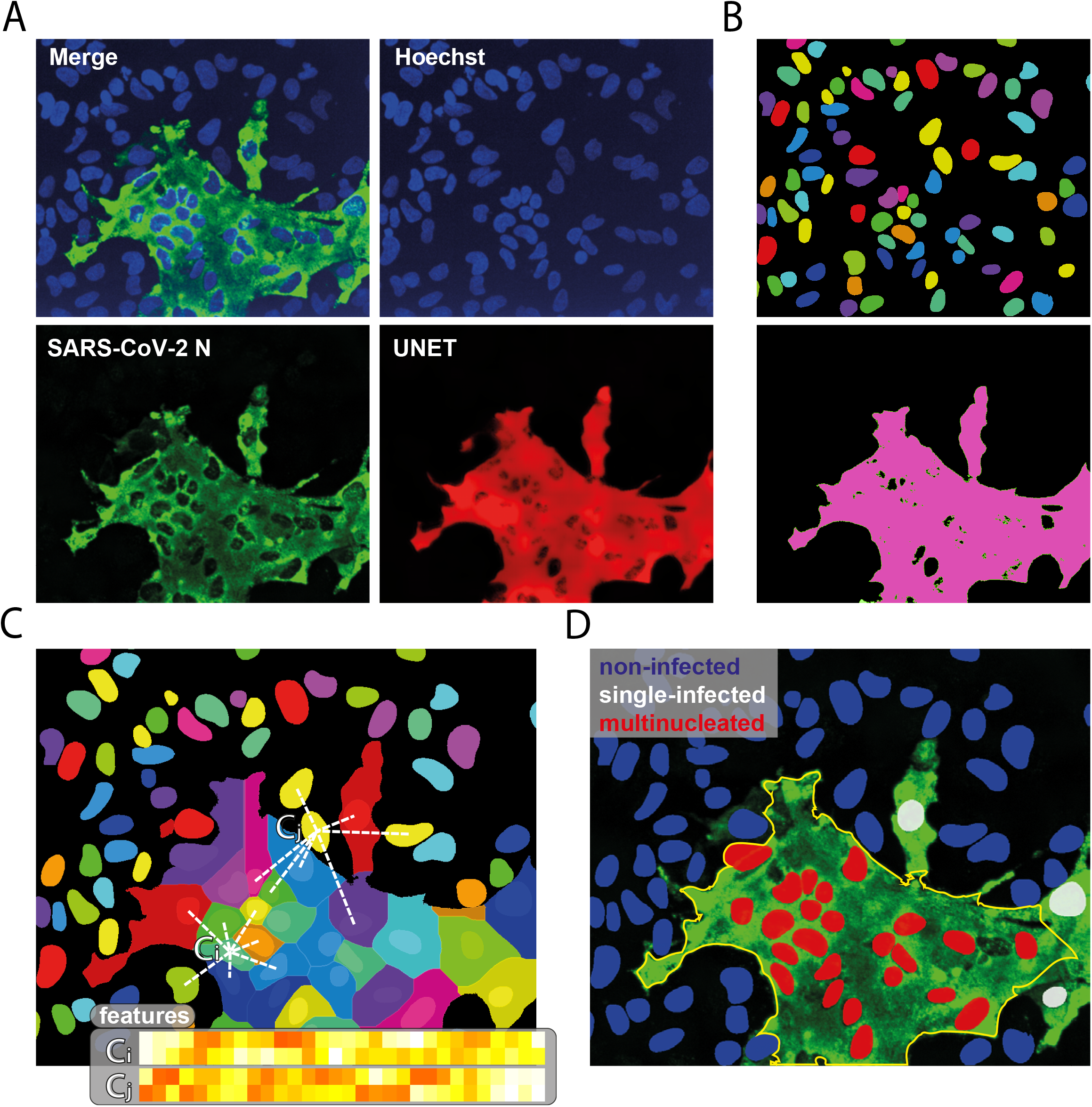
Image processing and phenotyping of SARS-CoV-2 infected cells. a) Original image of SARS-CoV-2 N signal (green) and enhanced image (red) using UNET deep learning algorithm. b) Single-cell segmentation of the nuclei using the nucleAIzer deep learning algorithm, and the cytoplasmic region based on global thresholding of the UNET enhanced image. c) Morphology, shape and intensity features of single-cells and their microenvironment are extracted. d) Machine learning-based phenotyping of single cells into non-infected, single-nuclei infected and multi-nucleated cells. Features include morphology, intensity and texture descriptor numbers. Ci: features of the i_th cell, Cj: features of the j_th cell.

**Extended Figure 3.**
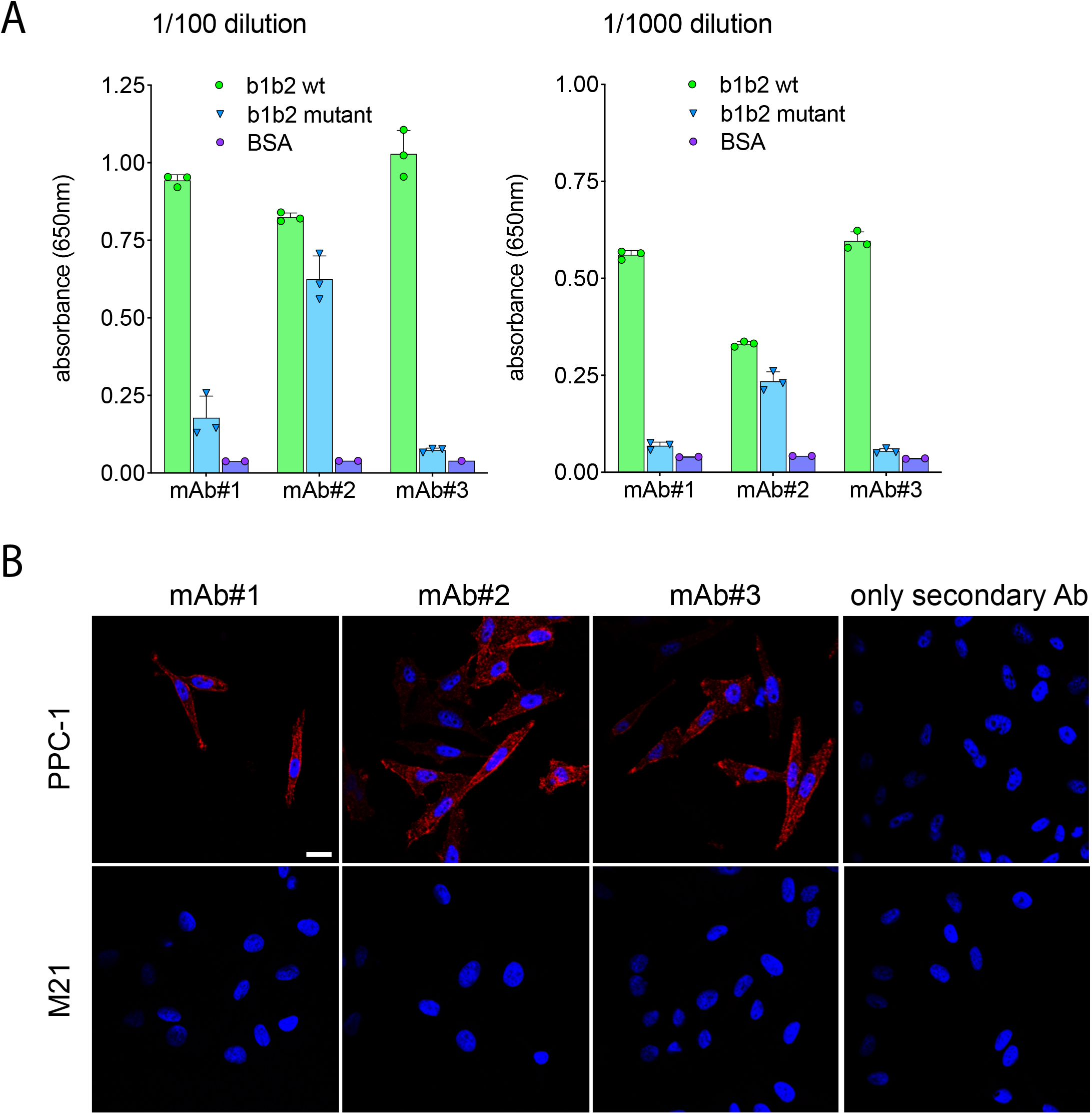
A) Binding of mAbs to wild type and mutant NRP-1. The binding, tested by ELISA at two different dilutions of mAbs, is represented as arbitrary units of absorbance. BSA was used as a negative control. B) Fluorescence confocal images of NRP1-positive prostate cancer cells (PPC-1) and NRP1-negative melanoma cells (M21) incubated with the monoclonal antibodies. Fixed cells on coverslips were incubated with the mAbs for 1 h and immunostained with the secondary antibody AlexaFluor 546 goat anti-mouse IgG. Representative images from three independent experiments are shown. Blue: DAPI (nuclei staining); Red: antibody signal. Scale bar; 20 μm.

